# Extracellular microRNA 3’ end modification across diverse body fluids

**DOI:** 10.1101/2020.03.24.006551

**Authors:** Kikuye Koyano, Jae Hoon Bahn, Xinshu Xiao

## Abstract

microRNAs (miRNAs) are small non-coding RNAs that play critical roles in gene regulation. The presence of miRNAs in extracellular biofluids is increasingly recognized. However, most previous characterization of extracellular miRNAs focused on their overall expression levels. Alternative sequence isoforms and modifications of miRNAs were rarely considered in the extracellular space. Here, we developed a highly accurate bioinformatic method, called miNTA, to identify 3’ non-templated additions (NTAs) of miRNAs using small RNA-sequencing data. Using miNTA, we conducted an in-depth analysis of miRNA 3’ NTA profiles in 1047 extracellular RNA-sequencing data sets of 4 types of biofluids. This analysis identified abundant 3’ uridylation and adenylation of miRNAs, with an estimated false discovery rate of <5%. Strikingly, we found that 3’ uridylation levels enabled segregation of different types of biofluids, more effectively than overall miRNA expression levels. This observation suggests that 3’ NTA levels possess fluid-specific information insensitive to batch effects. In addition, we observed that extracellular miRNAs with 3’ uridylations are enriched in processes related to angiogenesis, apoptosis and inflammatory response, and this type of modification may stabilize base-pairing between miRNAs and their target genes. Together, our study provides a comprehensive landscape of miRNA NTAs in human biofluids, which paves way for further biomarker discoveries. The insights generated in our work built a foundation for future functional, mechanistic and translational discoveries.

## INTRODUCTION

Recent studies revealed the existence of extracellular RNAs (exRNAs) in many types of biofluids^1–3^. exRNAs are mostly packaged in small extracellular vesicles, microvesicles or in complex with lipoproteins or ribonucleoproteins^4^, which protect them from degradation by ribonucleases. exRNA expression could be highly cell type- or disease-specific^5–7^, thus affording potential values as disease biomarkers^8^. Importantly, the functional roles of exRNAs are also starting to unfold^9–12^. For example, several studies reported the involvement of exRNAs in cell-to-cell communication in the local tumor microenvironment^13–15^. Furthermore, exRNAs in extracellular vesicles secreted from glioblastoma stem cells were shown to induce angiogenesis by reprogramming brain endothelial cells^9^.

The most-often studied exRNAs are microRNAs (miRNAs), small 18-22nt noncoding RNAs that are potent regulators of mRNA and protein expression levels^16^. Most previous studies on extracellular miRNAs focused on interrogating their overall expression levels. Nonetheless, many miRNAs assume multiple sequence forms resulted from alternative miRNA processing or post-transcriptional modification^17^. Specifically, a well-known class of post-transcriptional modification of miRNAs is non-templated addition (NTA)^18^. Two types of 3’ miRNA NTAs have been reported^19,20^, 3’ adenylation catalyzed by GLD2 and 3’ uridylation by the terminal uridyltransferase-4 and 7 (TUT4/TUT7). Both types of 3’ NTAs may affect miRNA targeting, stability, or turnover^21–25^.

Thus far, only a small number of studies examined miRNA NTAs in the extracellular space. For example, a study using cultured human B cells examined 3’ NTAs of intracellular and extracellular exosomal miRNAs^26^. The authors observed that 3’ NTAs of miRNAs showed distinct patterns in the two compartments, with 3’ adenylation more enriched intracellularly and 3’ uridylation overrepresented in exosomes. Another study examined global miRNA expression in blood cells, serum and exosomes^27^. They showed that 3’ NTA patterns clustered in a blood-lineage specific manner and extracellular 3’ NTAs were distinct from the intracellular profiles. These findings suggest that 3’ NTA patterns of miRNAs may carry specific information that segregates extra- and intracellular miRNA profiles. The mechanisms underlying this specificity remain unclear.

In this study, we developed a new analysis pipeline, called miNTA, to identify NTAs of miRNAs in any small RNA-sequencing (RNA-seq) data set and applied it to 1047 extracellular samples derived from 4 types of biofluids. To our best knowledge, this is the largest study of extracellular miRNA NTA profiles in biofluids. Although many studies have examined NTAs of intracellular miRNAs, the bioinformatic pipelines employed in most studies could be improved to enhance accuracy. Incorporating a number of stringency measures, our method achieves a low false discovery rate < 5%. Applied to the large number of biofluid samples, we observed 3’ uridylation and adenylation as the two most prominent types of 3’ NTAs in extracellular miRNAs, with the former being more prevalent than the latter. Our analysis showed that 5’ NTAs are unlikely present in miRNAs, or extremely rare if exist at all. The levels of 3’ NTAs varied widely across miRNAs. Importantly, 3’ NTA levels can be used to segregate different types of biofluids, more effectively than miRNA expression levels. We also observed that extracellular miRNAs with 3’ uridylations are enriched in processes related to angiogenesis, apoptosis and inflammatory response, and this type of modification may stabilize base-pairing between miRNAs and their target genes. Overall, our study provides global insights regarding 3’ end modifications of miRNAs in extracellular fluids.

## RESULTS

### miNTA: A bioinformatic pipeline to identify miRNA NTAs

To explore the diversity of NTAs in the extracellular space, we first developed a rigorous pipeline, called miNTA, to accurately identify NTAs of miRNAs in small RNA-seq data (Fig. 1A). While other methods to identify miRNA sequence variations exist^28–36^, our method aims to improve the mapping strategies and reduce false positive modifications (Methods). Briefly, in the mapping steps, we sequentially trim the ends of unmapped reads (separately for 3’ and 5’ ends) and check for one or more consecutive NTAs. To remove likely sequencing errors, we only consider candidate NTA sites with relatively high sequencing quality scores (PHRED ≥ 30). In addition, since miRNA ends can be heterogeneous and noisy^37,38^, we defined canonical isoforms of each miRNA in each sample and identified NTAs relative to the canonical 3’ or 5’ ends (Methods). These quality control steps ensure the accuracy of NTA predictions.

**Fig. 1.**
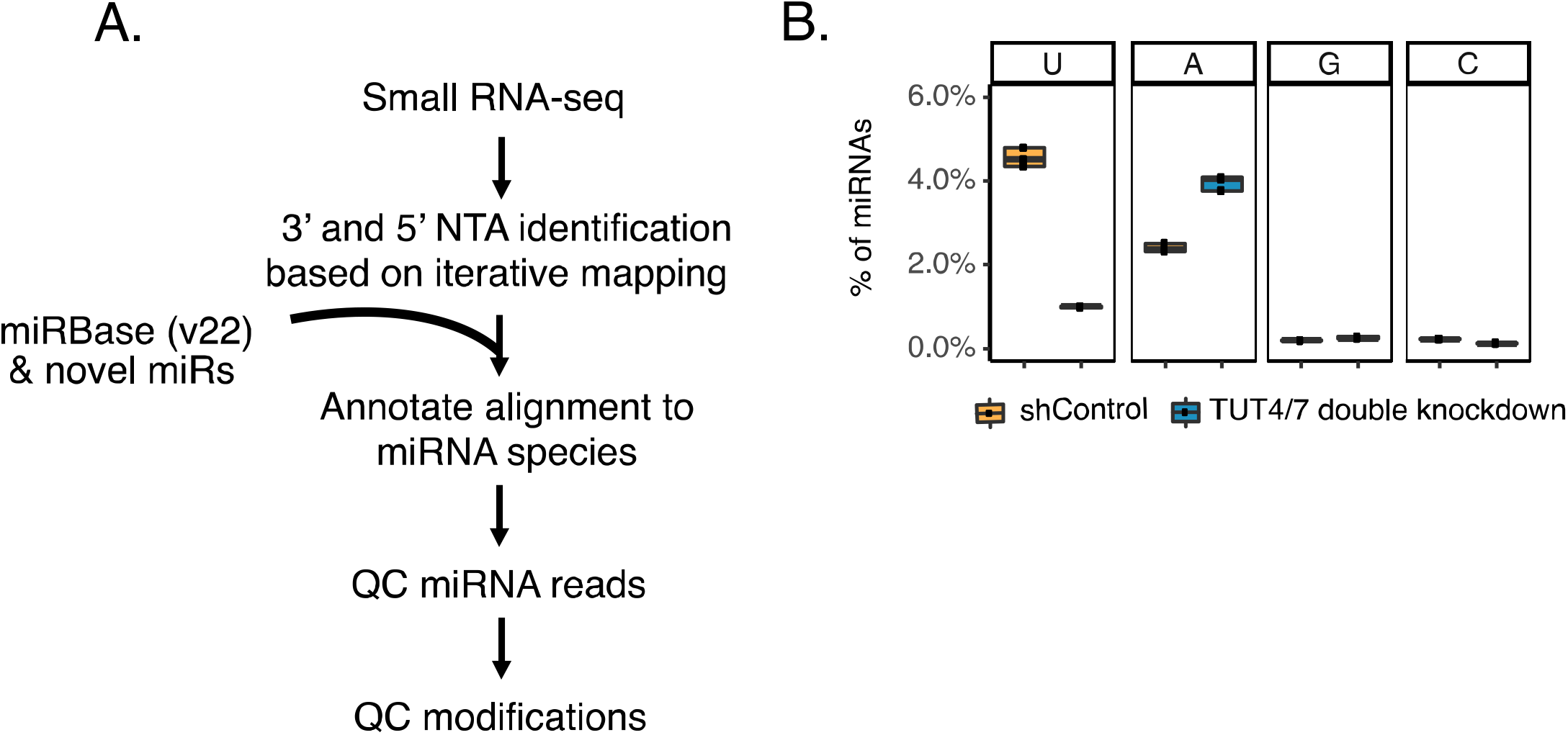
Identification of non-templated additions (NTAs) of miRNAs. (A) miNTA, a bioinformatic pipeline to identify miRNA NTAs. (B) Percentage of miRNAs with 3’ end non-templated mono-uridylation (U), adenylation (A), guanidylation (G) and cytidylation (C) identified by miNTA using small RNA-seq data derived from control (shControl) and TUT4/7 double KD HEK293 cells.

miRNA 3’ adenylation and uridylation are known to be mediated by specific enzymes such as, but not limited to, GLD-2 and TUT4/TUT7, respectively^16^. To evaluate if our pipeline detects biologically relevant NTAs, we performed double knockdown (KD) of the TUT4 and TUT7 enzymes in HEK293 cells, followed by small RNA-seq (Supplementary Fig. S1A). As expected, we observed reduced global 3’ uridylation levels of miRNAs upon TUT4/7 KD relative to the controls (Fig. 1B). In addition, a concomitant increase of 3’ adenylation levels was observed, consistent with previous literature^20,24^. These results confirm the validity of our pipeline in identifying biologically relevant NTAs.

### Comprehensive catalog of miRNAs in the extracellular space

To investigate the landscape of extracellular miRNA 3’ NTAs, we obtained small RNA-seq data sets (50 nt, strand-specific) of four bodily fluids of healthy subjects obtained from previous studies^39–43^ (Fig. 2A). In total, we analyzed 1047 data sets, including 399 plasma samples, 163 samples of small extracellular vesicles isolated from plasma, 167 serum, 69 cerebral spinal fluid (CSF), and 249 urine samples. As a comparison, we also analyzed 297 intracellular data sets of seven types of human peripheral blood cells (NK cells, B lymphocytes, cytotoxic T lymphocytes, T-helper cells, monocytes, neutrophils and erythrocytes) sorted from whole blood^27^. Following read mapping, we removed samples with <50,000 total reads mapped to miRNAs to ensure at least a modest sequencing coverage.

**Fig. 2.**
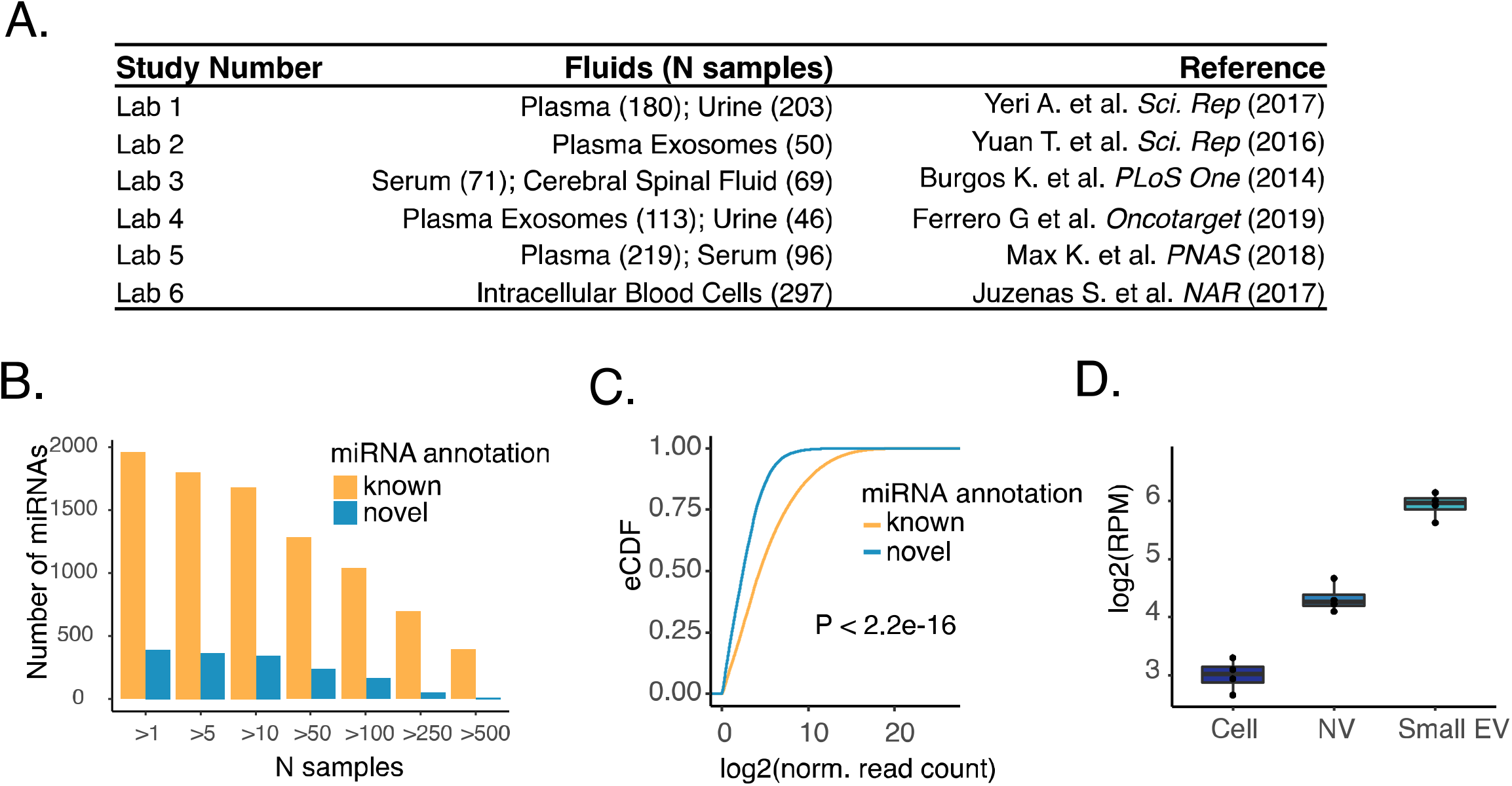
Generation of a comprehensive catalog of extracellular miRNAs. (A) Extracellular and intracellular small RNA-seq datasets used in this study. (B) Number of known and novel miRNAs observed in greater than N samples (x axis) across all data sets in (A). (C) Empirical cumulative distribution function (eCDF) of the abundance of known or novel miRNAs in all data sets in (A). Normalized read counts were calculated using DESeq2 (Methods). P value was calculated via a two-sided Kolmogorov–Smirnov (KS) test. (D) Expression of a novel miRNA (chr7_40460) in whole-cell lysates (Cell), non-vesicle extracellular (NV) and small extracellular vesicle (Small EV) fractions isolated from Gli36 cells.

Although intracellular miRNAs have been studied extensively, the repertoire of extracellular miRNAs is still being explored. To create a comprehensive list of human miRNAs, as the first step of the pipeline, we ran miRDeep2^44^ on all extracellular and intracellular data sets to identify novel miRNAs. This procedure identified a total of 404 novel miRNAs, present in more than one sample (Fig. 2B). Interestingly, 85.3% of these novel miRNAs were detected in more than 10 samples, a slightly higher percentage than that (81.7%) of known miRNAs. Nonetheless, novel miRNAs had relatively lower expression levels than known miRNAs (Fig. 2C), likely explaining their absence in the miRBase annotation. Notably, certain novel miRNAs may have higher expression levels in the extracellular space, such as the example shown in Fig. 2D (derived from paired intra- and extracellular data sets^45^, also see Supplementary Fig. S2). Henceforth, we include both annotated and novel miRNAs in the analysis.

### miRNA NTAs in the extracellular fluids and evaluation of the NTA pipeline

Next, we examined NTA profiles identified in individual miRNA reads, without grouping reads per miRNA. On average across fluids, 5.2% or 0.3% of all miRNA reads had 3’ or 5’ end NTAs, respectively (Fig. 3A). For intracellular samples, 11.2% or 0.2% of total miRNA reads had 3’ or 5’ end NTAs, respectively (Supplementary Fig. S3A).

**Fig. 3.**
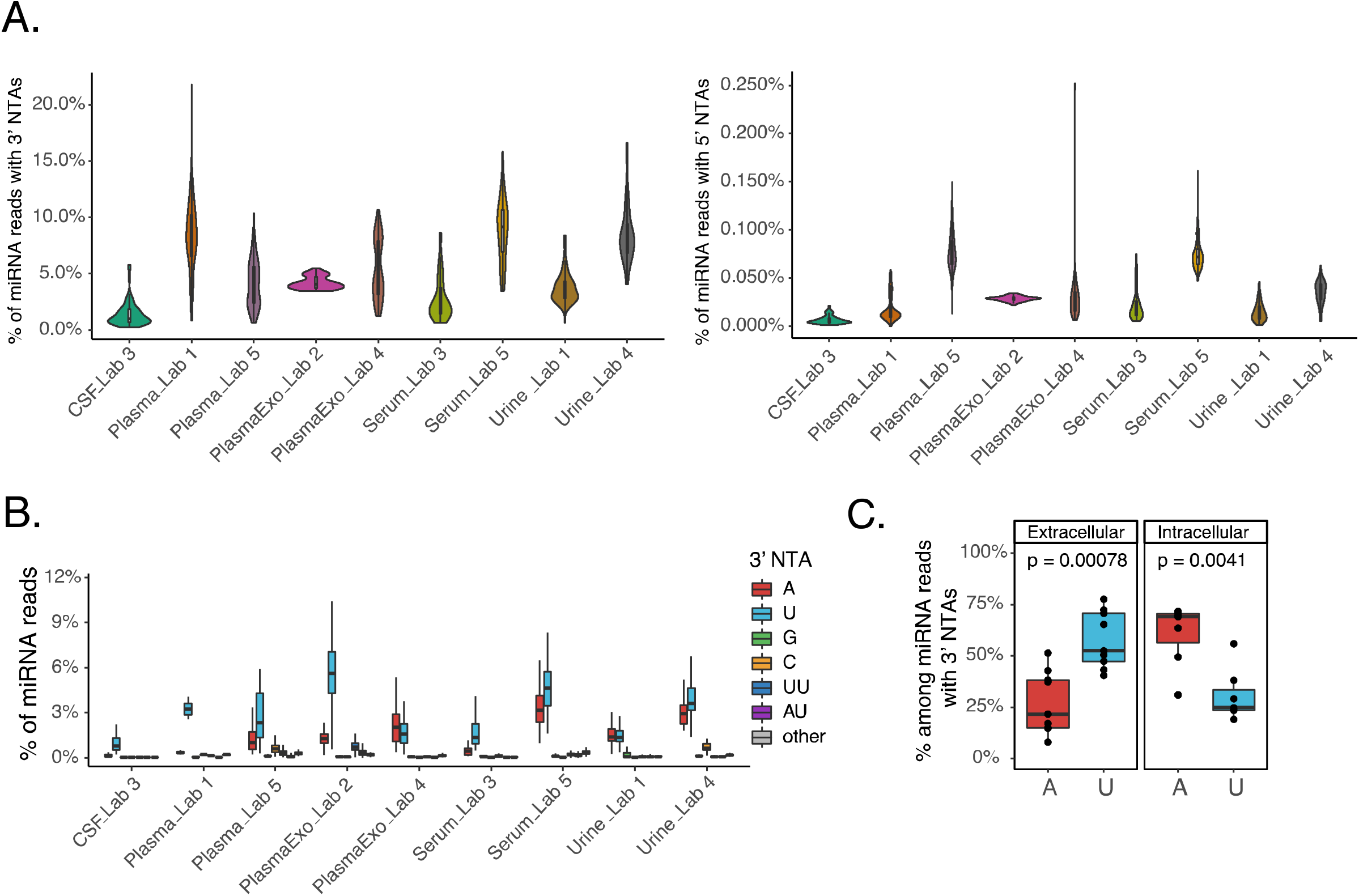
NTA profiles of extracellular miRNAs across biofluids. (A) Percentage of miRNA reads with 3’ (left) and 5’ (right) NTAs in each data set. (B) Nucleotide compositions of 3’ NTAs in miRNAs of each data set. (C) Average percentage of reads with 3’ uridylation or adenylation among all miRNA reads with 3’ NTAs. Each dot represents this average value for an extracellular fluid type or intracellular cell type in each study. P values were calculated via Wilcoxon rank-sum test.

We next examined the nucleotide composition of the 3’ NTAs across miRNA reads for each fluid. This analysis also allows us to evaluate the quality of our pipeline as miRNAs are expected to have predominantly two types of 3’ NTAs (adenylation and uridylation), based on previous reports^19,22,26,46^. On average, >94% of all mono 3’ end modifications in reads of extracellular samples were identified as either adenylation or uridylation, whereas 3’ end cytosine or guanosine additions were each less than 3.4% (Fig. 3B). Similar results were also observed for intracellular groups (Supplementary Fig. S3B). If the observed 3’ G or 3’ C modifications were assumed to be false positives, then the false discovery rate (FDR) of our predicted mono 3’ NTA-containing reads would be estimated to be <5% for exRNAs, and <2% for intracellular miRNAs. Note that these FDRs may be over-estimated since 3’ G or C NTAs may exist, although no known enzymes have been reported for these modifications in mammals.

Our bioinformatic pipeline also allowed an investigation of 5’ NTAs of miRNAs. In general, the prevalence of 5’ NTAs were much lower than that of 3’ NTAs (Fig. 3A). Despite this low level, a strong enrichment of cytosine among the 5’ NTAs was observed (Supplementary Fig. S4A). A low range of 5’ Cs were observed for 78 miRNAs in Plasma from Lab 5 (Supplementary Fig. S4B). However, the average level of 5’ C in miRNAs with this modification was very low (<0.028%). Since no known mechanisms exist to account for 5’ C modification of miRNAs in mammals, the observed 5’ C NTA may reflect technical rather than biological mechanisms. For example, the 5’ adaptor used in small RNA library generation ended with a C nucleotide, which may have been read as the first base of the read as a type of sequencing error. Another report also observed a high proportion of 5’ addition of C^34^. Although the 5’ C may be an experimental artifact, the fact that this strong nucleotide bias was observed despite the overall low prevalence of 5’ NTAs strongly supports the effectiveness of our pipeline.

### miRNA 3’ uridylation is relatively more prevalent than 3’ adenylation in biofluids

Next, we examined the landscape of 3’ NTAs of miRNAs by grouping reads for each miRNA. In this analysis (and all others henceforth), we required ≥2 reads in each sample carrying the NTA nucleotide, with ≥5 samples satisfying this requirement in the same data set. When summarizing results for each fluid data set, we further required that ≥10 total reads mapped to the corresponding miRNA in each sample.

Using the above criteria, we observed that 32% (496/1541) or 26% (405/1541) of all detected miRNA species had 3’ uridylation or 3’ adenylation, respectively, considering all the biofluid data sets. Although the total numbers of modified miRNA species were similar, 3’ uridylation in reads mapped to miRNAs was significantly more frequent than 3’ adenylation, 58% and 27% among all types of 3’ NTAs, respectively (Fig. 3C). In contrast, the opposite trend was observed in intracellular samples, where a larger fraction of miRNAs had 3’ adenylations than uridylations (61% and 31%, respectively) (Fig. 3C). Overall, miRNA 3’ uridylation was more frequent in extracellular data sets (58%) compared to intracellular data sets (31%) (Wilcoxon rank sums test, p value = 0.003).

### miRNAs exhibit a wide range of NTA levels in biofluids

The 3’ uridylation levels varied greatly across miRNAs in each type of fluid (Fig. 4A). While many miRNAs had relatively low 3’ U levels, a number of them had modest to high levels. For example, 39% (122/312) of 3’ U-harboring miRNAs in plasma (Lab 1 data set) had ≥20% 3’ U levels. In particular, the miRNA hsa-let-7f-2-3p demonstrated a high level of 3’ U (ranging 14-93%) across all extracellular data sets. A few top miRNAs with high 3’ U levels in different fluids are listed in Table 1 and Supplementary Table S1A. Note that the 3’ U levels of the same miRNA between different data sets may not be comparable due to the distinct experimental protocols used to generate each data set. Among the three studies where at least 10 novel miRNAs were detected with 3’ uridylation, two showed higher 3’ U levels in novel miRNAs than known ones (Wilcoxon rank-based test, Supplementary Fig. S5).

**Fig. 4.**
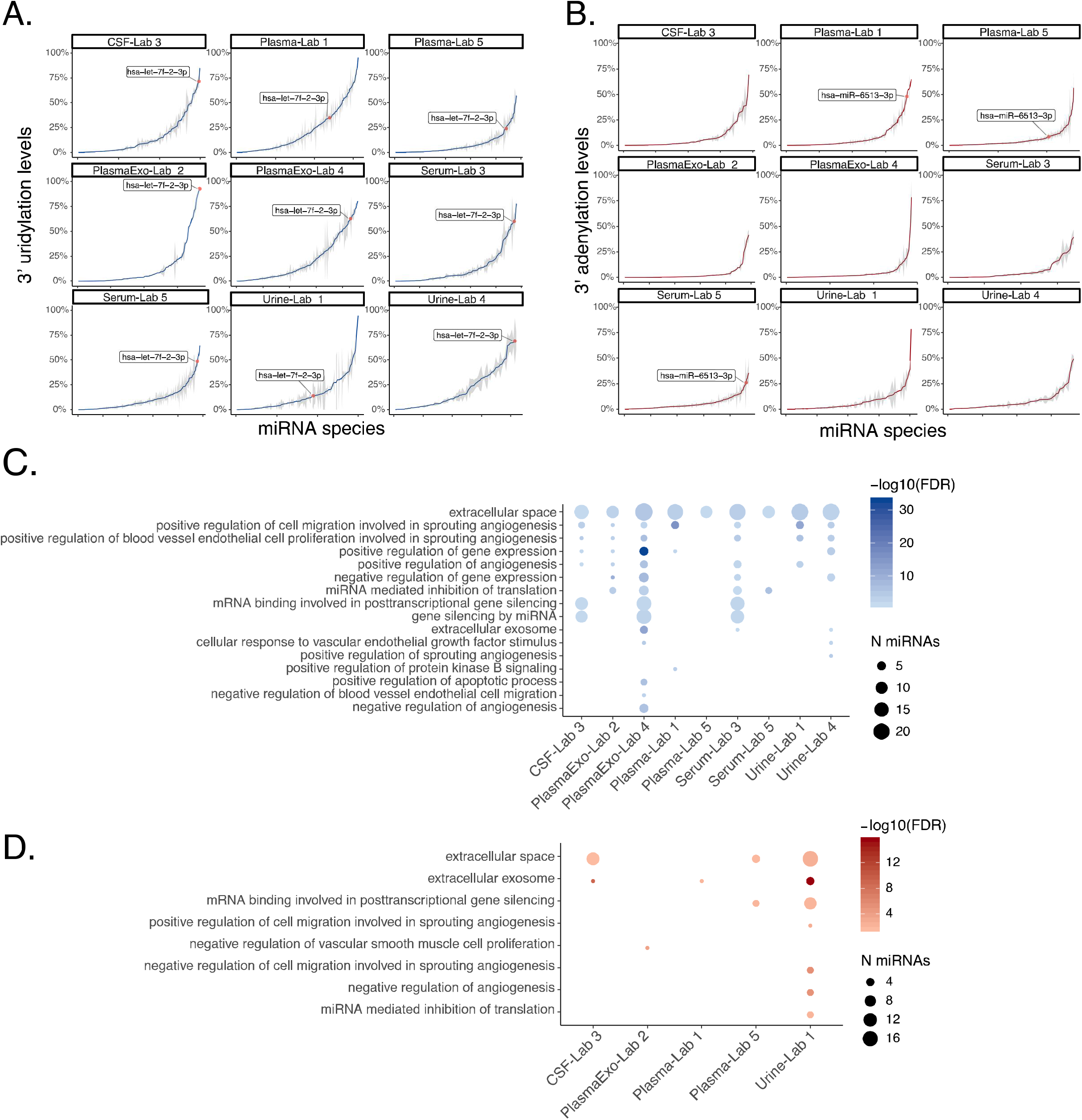
Characteristics of miRNAs with 3’ NTAs. (A) 3’ uridylation levels of miRNAs in each data set. Blue curves represent the average 3’ NTA levels and grey shades represent standard errors. The miRNA hsa-let-7f-2-3p is highlighted. (B) Similar to (A) but for miRNA 3’ adenylation levels. The miRNA hsa-miR-6513-3p is highlighted. (C) Gene ontology terms enriched among miRNAs with an average 3’ uridylation level ≥ 5% (FDR < 0.05, Methods). The size of the dots reflects the number of miRNAs in each GO term. (D) Similar to (C) but for miRNA 3’ adenylation.

**Table 1.**
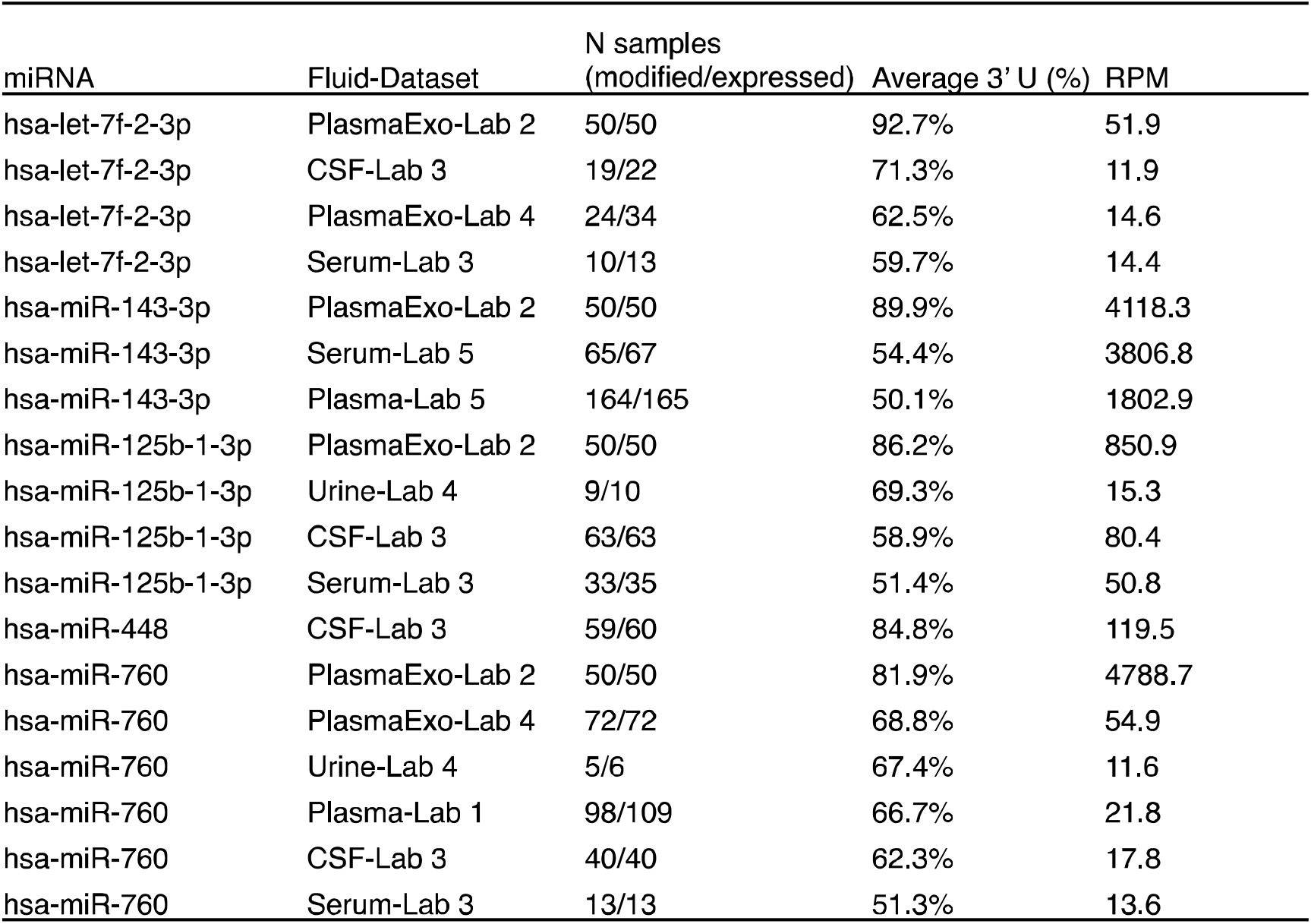
Top 5 3’ uridylated miRNA across biofluids. miRNAs were required to be modified and expressed (read count ≥ 10) in at least 5 samples from the original data set. Other data sets harboring the modified miRNA with an average 3’ uridylation level ≥ 50% are displayed. See Supplementary Table S1A for all miRNA 3’ uridylation levels.

Similarly, the 3’ adenylation levels also varied across miRNAs (Fig. 4B). Consistent with the previous observation of lower 3’ A levels compared to those of 3’ U, only 14.7% (27/189) of 3’ A-harboring miRNAs had a 3’ A level ≥20% in the plasma (Lab 1) data set. Nonetheless, a small number of miRNAs had considerable levels of 3’ adenylation (Table 2 and Supplementary Table S1B). An example is hsa-miR-6513-3p that demonstrated a high level of 3’ A (ranging 8-48%) across 3 extracellular data sets. No studies were detected with at least 10 novel 3’ adenylated miRNAs.

**Table 2.** Top 5 3’ adenylated miRNA across bioflui ds. miRNAs were required to be modified and expressed (read count ≥10) in at least 5 samples from the original data set. Other data sets harboring the modified miRNA with an average of 3’ adenylation level ≥ 50% were displayed. See Supplementary Table S1B for all miRNA 3’ adenylation levels.

| miRNA | Fluid-Dataset | N samples<br>(modified/expressed) | Average 3' A (%) | RPM |
| --- | --- | --- | --- | --- |
| hsa-miR-944 | CSF-Lab 3 | 7/7 | 69.2% | 9.5 |
| hsa-miR-1237-3p | Plasma-Lab 5 | 5/6 | 56.4% | 3.3 |
| hsa-miR-1237-3p | Plasma-Lab 1 | 6/6 | 44.3% | 4.8 |
| hsa-miR-6775-3p | Plasma-Lab 1 | 18/20 | 54.7% | 7.8 |
| chr2_27505 | PlasmaExo-Lab 4 | 6/7 | 52.4% | 5.0 |
| hsa-miR-205-5p | Urine-Lab 4 | 22/22 | 49.4% | 295.8 |
| hsa-miR-205-5p | PlasmaExo-Lab 4 | 6/9 | 21.0% | 34.5 |

### NTAs often occur in miRNAs relevant to angiogenesis or signaling

To gain insights on the functional relevance of 3’ uridylation of miRNAs, for each fluid in each data set, we performed Gene Ontology (GO) enrichment analysis on miRNAs that have an average 3’ uridylation level ≥ 5% in at least 50% of samples. As background controls, miRNAs without 3’ modifications in our data were chosen randomly by matching their expression levels with those that harbored 3’ Us (Methods). Interestingly, the GO term “extracellular space” was significantly enriched (FDR < 0.05) for all data sets (Fig. 4C). Since this analysis controlled for expression, this observation supports the enrichment of 3’ uridylation of miRNAs in the extracellular space. In addition, we observed terms related to angiogenesis, apoptosis, gene regulation and inflammatory response, indicating the involvement of 3’ U-containing miRNAs in these processes. Since miRNAs in the let-7 miRNA family are known to be enriched with 3’ uridylation^47^, we repeated this analysis by excluding let-7 miRNAs. Similar enriched GO terms were observed, supporting the generality of the results for diverse miRNA species (Supplementary Fig. S6). For 3’ adenylated miRNAs, an enrichment for angiogenesis-related terms was observed, but mostly from the urine data set (Lab 1) (Fig. 4D).

### 3’ uridylation levels of miRNAs segregate biofluids robust to batch effects

Given the wide range of 3’ uridylation levels in miRNAs, we asked whether this modification can help to segregate different types of fluids. tSNE clustering on 3’ uridylation levels of all expressed miRNAs showed that samples derived from similar fluid types tend to cluster together (Fig. 5A). Specifically, serum, urine and plasma samples each contained data sets generated by two different labs. For these fluids (especially serum and urine), we observed that samples of the same fluid type generated by different labs largely clustered together. This observation strongly suggests that 3’ uridylation levels are informative in segregating fluid types. In addition, we observed that intracellular blood cell types descending from the lymphoid lineage, such as T lymphocytes (CD4+, CD8+), B lymphocytes (CD19+) and natural killer cells (CD56+), clustered separately from cell types of the myeloid lineage. Myeloid cell types such as erythrocytes (CD235a), neutrophils (CD15+), and monocytes (CD14+) also clustered separately from each other. Cell types clustering by blood lineage based on their 3’-end composition was previously reported^27^. Exosomal plasma samples did not cluster with plasma/serum samples, likely due to batch effects or the biological difference between exosomal and total extracellular miRNA contents. Segregation of samples based on 3’ adenylation levels was not as effective as using 3’ uridylation levels (Supplementary Fig. S7A), although serum and plasma samples from different labs did cluster together.

**Fig. 5.**
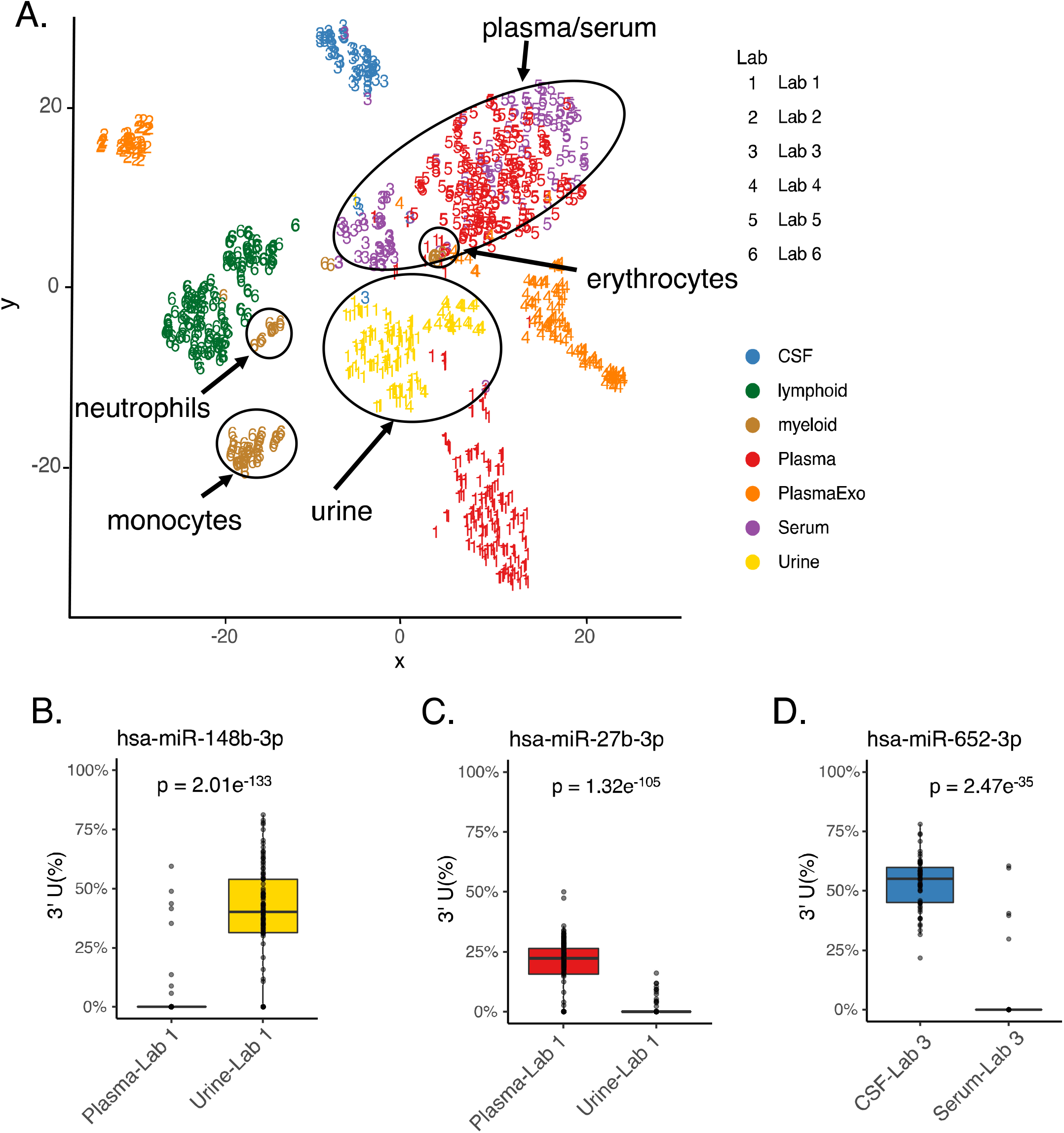
Distinct miRNA 3’ uridylation profiles across fluids. (A) tSNE clustering of samples using miRNA 3’ uridylation levels. miRNAs expressed with a minimum read count of 10 were included for this analysis. (B-D) Example miRNAs observed with differential 3’ uridylation levels between fluids of the same study. P values were calculated via REDITs (Methods).

As a comparison, we also performed tSNE on normalized miRNA expression values (Methods). This analysis showed exacerbated batch effects where data generated from the same lab largely clustered together, instead of clustering by fluid types (Supplementary Fig. S7B). PCA analysis of these samples showed improved fluid segregation, but still confounded by batch effects to some degree (Supplementary Fig. S7C). These results are consistent with observations made in previous studies where batch effects confounded the clustering of miRNA expression across samples^4^.

### Comparison of 3’ uridylation of miRNAs between biofluids

The above results suggest that different types of biofluids have distinct levels of 3’ uridylation. Thus, 3’ NTAs add another layer of information to distinguish fluid types. Nonetheless, since batch effects may not be completely absent, we avoided carrying out direct comparisons of the quantitative levels of NTAs across different studies. Instead, we performed differential modification analysis between fluids of the same study (Methods^48^).

For this analysis, we included miRNAs expressed in at least 20 samples in both fluids of a study. Overall, we identified 62 miRNAs with differential 3’ uridylation levels (≥5% difference in modification levels, FDR<0.05) (Supplementary Table S2). Specifically, 25 miRNAs had differential 3’ uridylation between urine and plasma (Lab 1), 11 between CSF and serum (Lab 3), 12 between urine and exosomes from plasma (Lab 4), and 29 between plasma and serum (Lab 5). Fig. 5B-D show three example miRNAs whose 3’ uridylation levels were high in one fluid, but almost zero in another.

### 3’ uridylation may increase miRNA base-pairing to its targets

Given the relatively high levels of 3’ uridylation in extracellular miRNAs, we hypothesized that this modification may affect the base-pairing between miRNAs and their targets. We focused on the top 20 unique miRNAs with highest average 3’ uridylation levels (per fluid) across all samples (Methods). Using RNAhybrid^49^, we observed that 3’ uridylation tends to stabilize the interaction between miRNAs and their targets, to an extent much greater than 3’ NTAs of G, C or A nucleotides (Fig. 6A, Chi-squared test p < 0.05). All miRNAs were significant against at least one of the background nucleotides. Target genes that paired with the 3’ U of these miRNAs were enriched in a variety of processes including extracellular exosome, integral protein of plasma membrane and negative regulation of apoptosis (Fig. 6B). Thus, it is possible that 3’ uridylation of extracellular miRNAs may affect miRNA targeting once taken up by recipient cells.

**Fig. 6.**
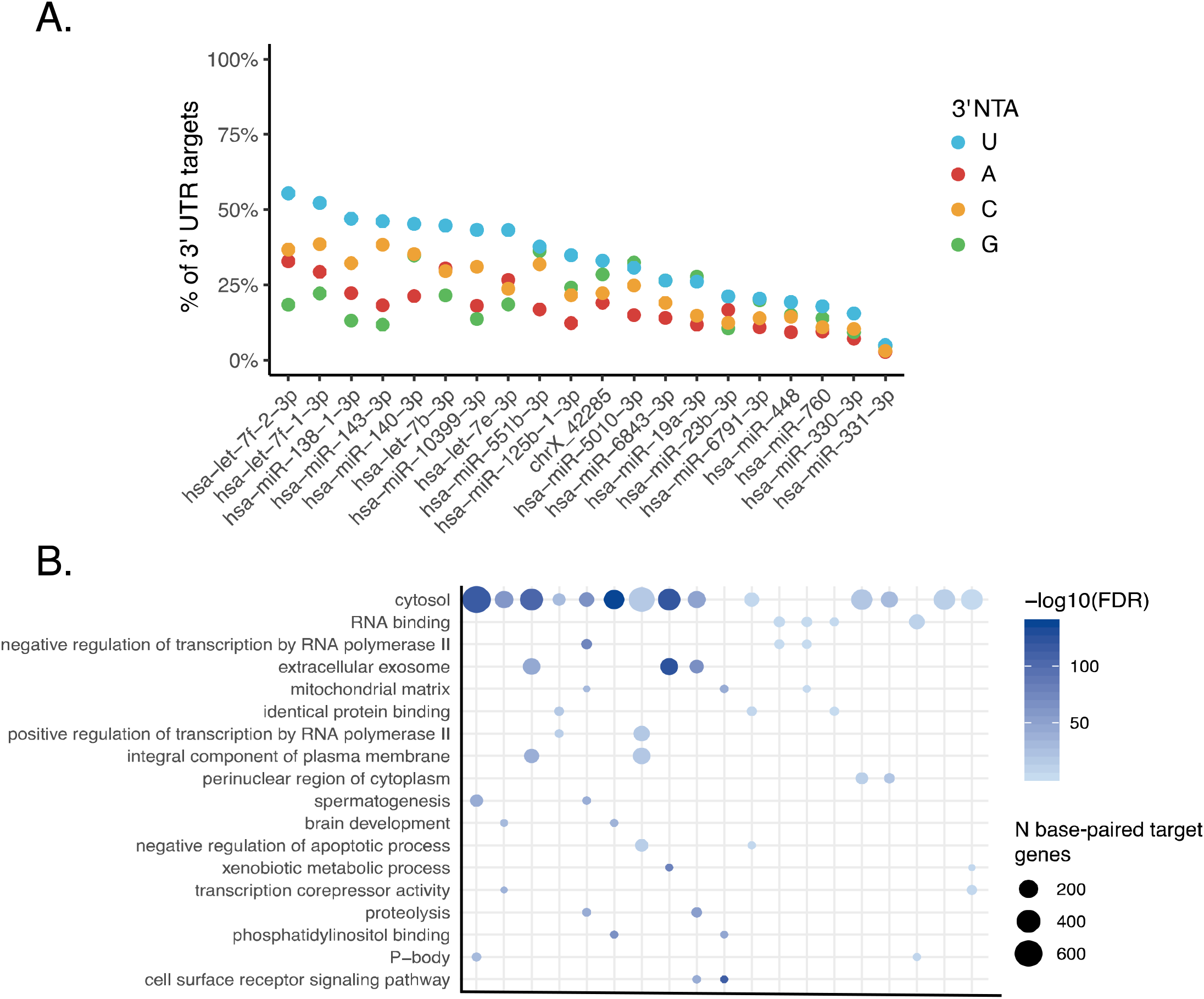
3’ U base-pairs with predicted miRNA targets more often than other 3’ NTAs. (A) Percentage of predicted miRNA 3’ UTR targets that base-pair with 3’ U (or A, C, G) for the top 20 unique miRNAs with highest average 3’ uridylation levels (per fluid) across all samples (Methods). For all miRNAs, the number of 3’ Us base-paired with predicted miRNA targets is higher than at least one background nucleotide (A, C or G) (Chi-squared test p <0.05). (B) For each miRNA, the top 5 enriched GO terms among its target genes that base-paired with the 3’ U were collected (Methods). Enriched terms observed in at least 2 data sets are shown (FDR < 0.05). The size of the dots represents the number (N) of base-paired target genes.

## DISCUSSION

We developed miNTA, a new bioinformatic pipeline, to identify 3’ NTAs of miRNAs. With different types of data sets, we demonstrated that the pipeline yields accurate and sensitive predictions. Using miNTA, we analyzed the global landscape of 3’ NTAs of extracellular miRNAs in >1000 biofluid samples, the largest study so far for extracellular miRNA modifications (to the best of our knowledge). We made a number of notable observations: (1) 3’ uridylation levels of miRNAs are higher in the extracellular space than 3’ adenylation levels, whereas the opposite was observed for intracellular miRNAs. (2) The level of 3’ NTAs varied widely across miRNAs, with some miRNAs demonstrating nearly 100% 3’ uridylation in certain biofluids. (3) 3’ uridylation levels of miRNAs can inform segregation of different types of biofluids. In addition, such segregation was more effective than that achieved by miRNA expression levels and largely robust to batch effects. (4) Extracellular miRNAs with 3’ uridylations are enriched in functional categories related to angiogenesis, apoptosis and inflammatory response, and 3’ uridylation may stabilize base-pairing between miRNAs and their target genes.

The effective segregation of biofluids by 3’ uridylation levels of miRNAs indicates that this type of miRNA modification possesses fluid-specific signatures. Thus, miRNA modification levels could serve as biomarkers that are less susceptible to batch effects. This property is likely due to the fact that modification levels are normalized metrics relative to the total miRNA expression. Differences in uridylation levels across fluids may be due to different cell types contributing distinct miRNA species into the extracellular space. Together, these results show that 3’ modification levels of miRNAs add an important layer of information to characterize extracellular RNA content.

The enrichment of 3’ uridylated extracellular miRNAs in angiogenesis, apoptosis and inflammatory responses suggests that this modification may have important functional relevance. For example, angiogenesis is a tightly regulated process that requires endothelial cells to be in close communication with their environment^50^. Extracellular vesicles were previously shown to play a role in regulating angiogenesis^9,50^. Thus, 3’ uridylation of miRNAs may be an important aspect contributing to this process. It should be noted that the functional enrichment analysis controlled for extracellular expression levels of 3’ uridylated miRNAs. Thus, the enriched categories reflect functions that are particularly relevant to 3’ uridylated miRNAs instead of extracellular miRNAs in general. We showed that 3’ uridylation may strengthen miRNA targeting. Interestingly, a previous study reported that 3’ uridylation of miR-27a induced target repression of ‘non-canonical’ sites with only partial seed-match and extensive 3’ end pairing^45^. Future studies are needed to understand the functional relevance of 3’ uridylation in extracellular miRNAs.

In summary, we presented an accurate pipeline to identify 3’ NTA patterns in extracellular miRNAs. Our large-scale analysis of data from different human biofluids supports that fluid-specific signatures exist in 3’ modifications of miRNAs. Our work extends the basis for future studies on the functional relevance of extracellular miRNA modifications and their values in biomarker discoveries.

## METHODS

### miNTA: read mapping

For each small RNA-seq data set, adapters and low-quality nucleotides were removed from raw fastq sequences using cutadapt (v.1.11). To enable comprehensive read mapping, miNTA includes a multi-step mapping strategy. First, all reads were aligned to the human genome (hg19) using Bowtie (version 1.1.2)^51^, allowing up to one mismatch and retaining uniquely mapped reads only. This stringency aims to minimize ambiguous mapping results. Next, the unmapped reads were trimmed by 1 nucleotide at their 3’ ends and realigned to the human genome according to the same requirements as described above. Unaligned reads from this step were trimmed again and aligned for another round. The above procedures were repeated using all original reads to carry out 5’ trimming and identify 5’ end NTAs. Following the above analysis, all remaining unmapped reads were restored to their original sequences and realigned after trimming 1 nucleotide each from the 3’ and 5’ end, respectively. Finally, the mapped reads (trimmed or untrimmed) were examined relative to the human genome reference to identify 3’ and 5’ NTAs.

### miNTA: quality control (QC) procedures

#### QC of mapped reads

Incorrect mapping can lead to false positive predictions of NTAs. To ensure accurate read mapping, we realigned reads that mapped to miRNAs using BLAT^52^ against the human genome. Those reads that did not yield consistent alignment results by BLAT and Bowtie were removed from further analysis.

#### Canonical end positions of miRNAs

It is known that miRNAs may express alternative isoforms (isomiRs). The existence of isomiRs, especially the low abundant ones, may complicate the identification of NTAs. Thus, we incorporated a procedure to identify canonical end positions of miRNAs. For each miRNA in each sample, the canonical 3’ end was defined as the position of the last nucleotide in the miRNA read sequence that matches the reference genome and supported by at least 50% of reads aligned to this miRNA. The canonical 5’ end of each miRNA was defined similarly for each sample. Subsequently, NTAs were identified relative to the canonical end positions for each miRNA.

#### QC of modifications

The predicted NTAs were further examined to eliminate those that may reflect genetic variants or technical artifacts. First, predicted NTAs that overlapped known SNPs or genetic mutations were removed^53–57^. Second, NTAs with a 100% modification level were removed as they may be due to mapping errors (similarly as in other studies of single-nucleotide variants^58,59^). Third, to minimize false positives due to likely sequencing errors, we removed reads with a PHRED score < 30 at the position corresponding to the NTA. Lastly, for NTAs of each miRNA, we required at least 2 reads with the modification and the NTAs observed in at least 5 samples within the same fluid and data set group.

### Novel miRNA prediction

Novel miRNAs were predicted using miRDeep2^44^ by combining reads from all samples of the same fluid (or cell type) in the same study. To obtain high confidence predictions, we imposed the following criteria, similar to previous studies^60^: (1) a miRDeep2 score > 0; (2) ≥ 5 reads mapped to the passenger strand and ≥10 reads to the primary strand. For overlapping predictions, the one with the highest miRDeep2 score was chosen. The final set of putative novel miRNAs were combined with known miRNA annotations (miRBase V22)^61^ for subsequent analyses.

### miRNA read count normalization across data sets

To obtain miRNA expression levels, DESeq2^62^ (version 1.14.1) was used to normalize miRNA read counts across data sets. miRNAs associated with at least 10 reads in at least 50% of all samples were used to generate the DESeq2 scale factor for normalization.

### Gene ontology enrichment analysis

Gene ontology (GO) terms for miRNAs were downloaded from http://geneontology.org/gene-associations/goa_human_rna.gaf.gz^63^. For each data set, miRNAs expressed in ≥ 50% of samples and with an average 3’ uridylation level of ≥5% were included for the GO analysis. For each query miRNA, a control miRNA without 3’ modifications was chosen randomly that matches the expression level of the query (± 20% relative to the query). GO terms present in the sets of query miRNAs and matched controls were collected, respectively. The process was repeated 10,000 times to construct 10,000 sets of control miRNAs, where each set has the same number of miRNAs as the query set. Query miRNAs that had less than 3 candidate controls were not included in this analysis. For each GO term associated with at least 2 query miRNAs, a Gaussian distribution was fit to the number of control miRNAs also associated with this term to calculate a p value. Significant GO terms were defined as those with FDR < 0.05.

### tSNE and PCA clustering

miRNAs expressed with a minimum read count of 10 were included for clustering analysis. tSNE and PCA clustering were performed using the package Rtsne and prcomp, respectively. The expression was set to 0 in samples where a miRNA has no reads. Levels of NTAs or Log2 of the DEseq2 normalized expression was used in these analyses.

### Differentially modified miRNA between fluids

Differential modification of miRNAs between two fluids was performed using the REDITs method^48^. miRNAs with an effect size ≥ 5% between fluids in expressed samples were included. Significant miRNAs were required to be expressed (read count ≥ 10) in at least 20 samples in both tested fluids with a FDR < 0.05.

### miRNA target analysis

RNAhybrid (version 2.1.2)^49^ with default settings was used to estimate the minimum free energy between miRNAs and putative target 3’ UTR sequences. RNAhybrid p value was required to be <0.05 to call a significant minimum free energy binding site. Putative sites that base-paired with different types of 3’ NTAs (A, U, C, G) were then examined. GO enrichment of target genes pairing with the 3’ U NTAs was performed similarly as described above. For each miRNA, control genes were chosen from those putative target genes without 3’ U-pairing and with matched gene length as the query genes (±10%). Each query gene was required to have at least 10 controls to be included in the analysis.

### Cell Culture

Human embryonic kidney cell line (HEK293T) was obtained from the ATCC. Cells were maintained in Dulbecco’s modified Eagle’s medium containing 10% fetal bovine serum (FBS) with antibiotics at 37C in 5% CO2.

### shTUT4 and shTUT7 knockdown

We used lentivirus-packaged short hairpin RNA (shRNA) to knock down TUT4 and TUT7. The shRNA sequences (purchased from IDT) were obtained from a previous study^20^. Co-transfection of pCMV-d8.91, pVSV-G and pLKO1 into HEK293T cells was performed using Lipofectamine 3000 (Thermo Fisher Scientific, Cat# L3000-008). Lentiviruses were collected from conditioned media 48hrs after transfection. Lentivirus-containing media was filtered using 0.45 μM PES syringe filter (VWR) and mixed with polybrene (8 μg/ml). Following 24hrs of infection, cells were incubated with puromycin (1 μg/ml) for 3-4 days. To make double knock down cell lines, second round of infected cells were incubated with hygromycin (200 μg/ml) for 3-4 days. Knockdown efficiency was evaluated by Western blot using TUT4 (Proteintech Inc, Cat# 18980-1-AP), TUT7 (Bethyl Laboratories cat# A305-089A) and beta-actin antibodies (Santa Cruz Biotech, Cat# sc-47778 HRP).

### Extracellular and intracellular RNA isolation

Lentivirus-infected HEK293 cells were washed three times with PBS and the medium was switched to serum-free medium containing antibiotic-antimycotic (Thermo Fisher Scientific, Cat# 15240112). Following 24hrs incubation, the cell culture medium was collected and centrifuged at 2,000 g for 15 min at room temperature. To thoroughly remove cellular debris, the supernatant was centrifuged again at 12,000 g for 35 min at room temperature. Then the conditioned medium was used for RNA extraction with Trizol (Thermo Fisher Scientific, Cat# 10296028). Intracellular RNA was isolated using Direct-zol RNA mini prep kit (Zymo Research) from the same culture dish for extracellular RNA.

### Small RNA library preparation

Small RNA sequencing libraries were generated using the NEBNext Small RNA library Prep kit and NEBNext multiplex oligos for Illumina according to the manufacturer’s instructions (New England Biolabs, E7300). The final small RNA libraries were purified from 6% PAGE gel, and their concentrations were measured by Qubit fluorometric assay (Life Technologies). Libraries were sequenced on an Illumina HiSeq-3000 (50-bp singleend).

## ACKNOWLEDGEMENTS

We thank members of the Xiao laboratory for helpful discussions and comments on this work. We acknowledge the data production efforts and the subject donors for making available the data sets used in this study. The Extracellular Small RNA Profiles in Plasma, Urine and Saliva from College Athletes Study (dbGAP phs001258) was made possible by the NIH Common Fund Program on Extracellular RNA Communication Grant UH3 TR000891, Riddell and BRG Sports, the Flinn Foundation grant awards #1994 and #2307, the staff and athletes at Arizona State University, and TGen faculty and interns. Thanks to The Michael J Fox Foundation for supporting the collection of dbGAP phs00727, the subject donors to the program at Sun Health Research Institute. This work was supported in part by grants from the National Institute of Health (U01HG009417, R01AG056476 to X.X) and the Jonsson Comprehensive Cancer Center at UCLA. K.K. was supported by a Eureka Scholarship from the Department of Integrative Biology and Physiology at UCLA.

