## Supplementary Figs for "Extracellular microRNA 3’ end modification across diverse body fluids"

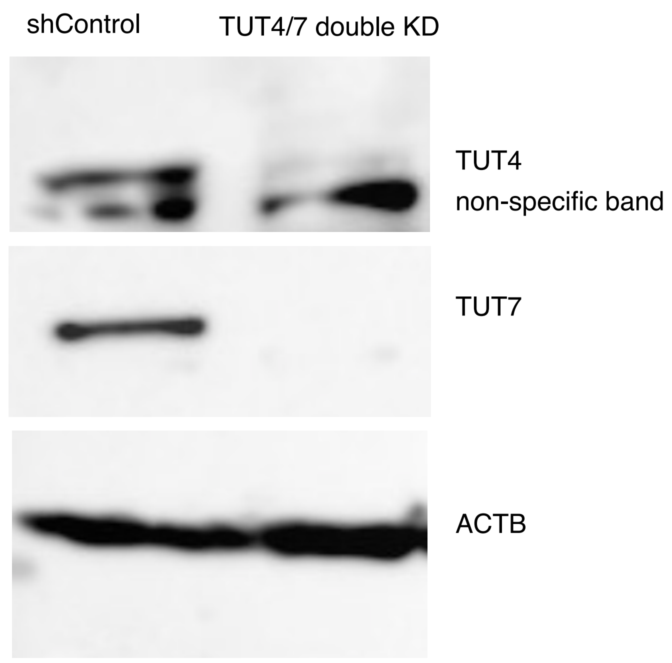


**Supplementary Fig. S1. TUT4/7 double knockdown to validate miNTA pipeline**
Western blot of control (shControl) and TUT4/7 double KD in HEK293 cells.

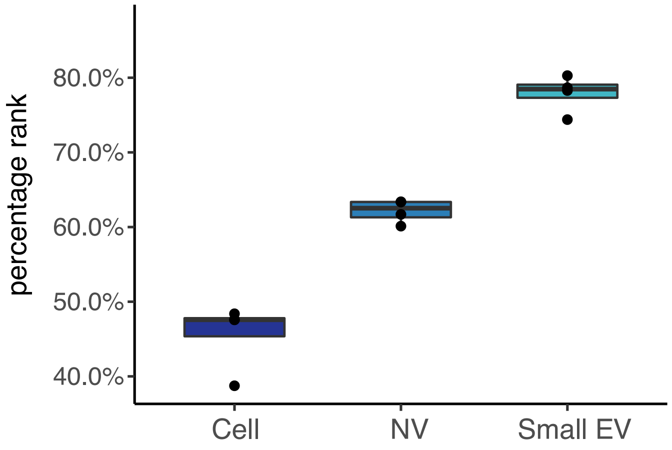


**Supplementary Fig. S2. A novel miRNA with higher expression levels in the extra-cellular space.** Percentage rank of a novel miRNA (chr7_40460) in whole-cell lysates (Cell), non-vesicle extracellular (NV) and small extracellular vesicle (small EV) fractions isolated from Gli36 cells.

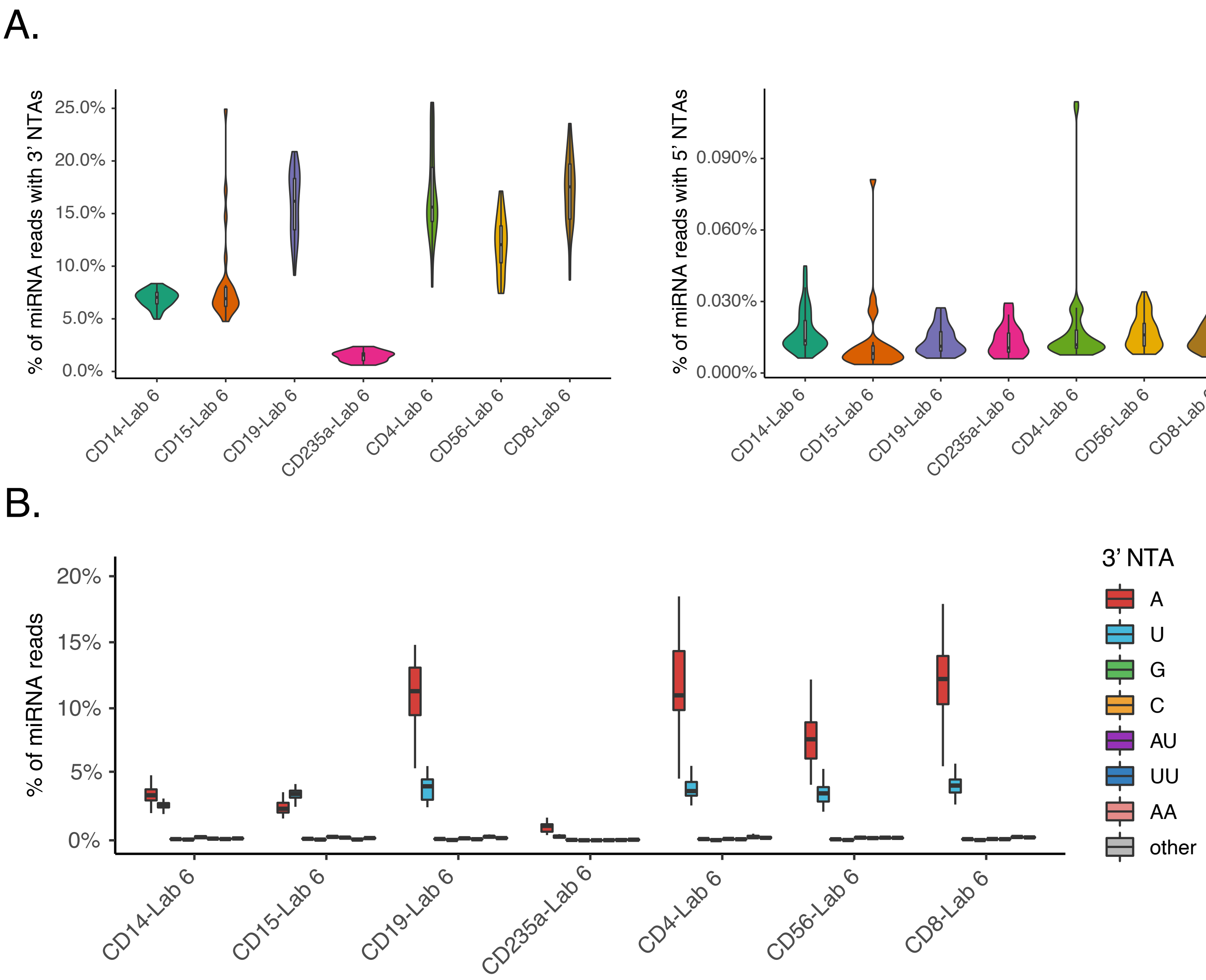


**Supplementary Fig. S3. NTA profiles of intracellular miRNAs across human peripheral blood cells.**

(A) Percentage of miRNA reads with 3' (left) and 5' (right) NTAs in each dataset.(B) Nucleotide composition of miRNAs with 3' NTAs across all samples in each intracellular blood-cell type.

A.

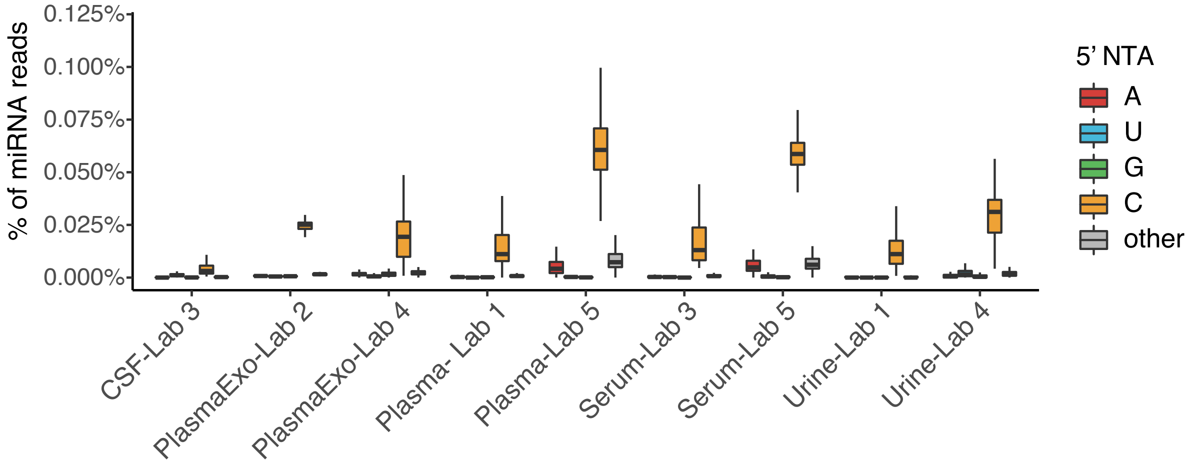


B.

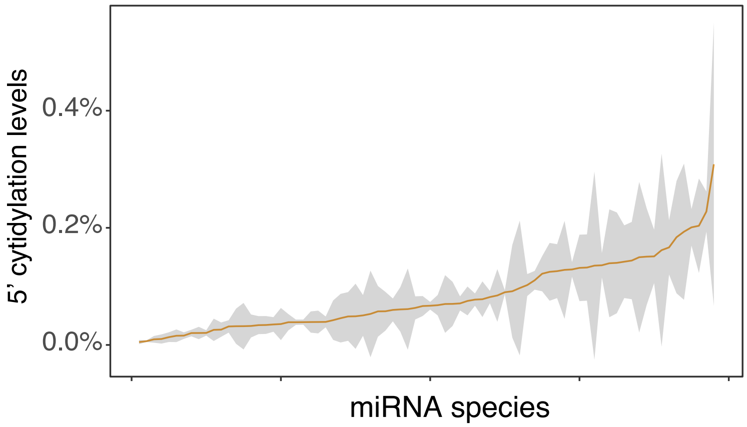


**Supplementary Fig. S4. miRNA 5' NTAs across biofluids.**
(A) Nucleotide composition of miRNAs with 5' NTAs across all samples in extracellular fluids from each study. (B) 5' cytidylation levels of miRNAs in Plasma-Lab 5. Orange curve represents the average 5' NTA levels and grey shades represent standard errors.

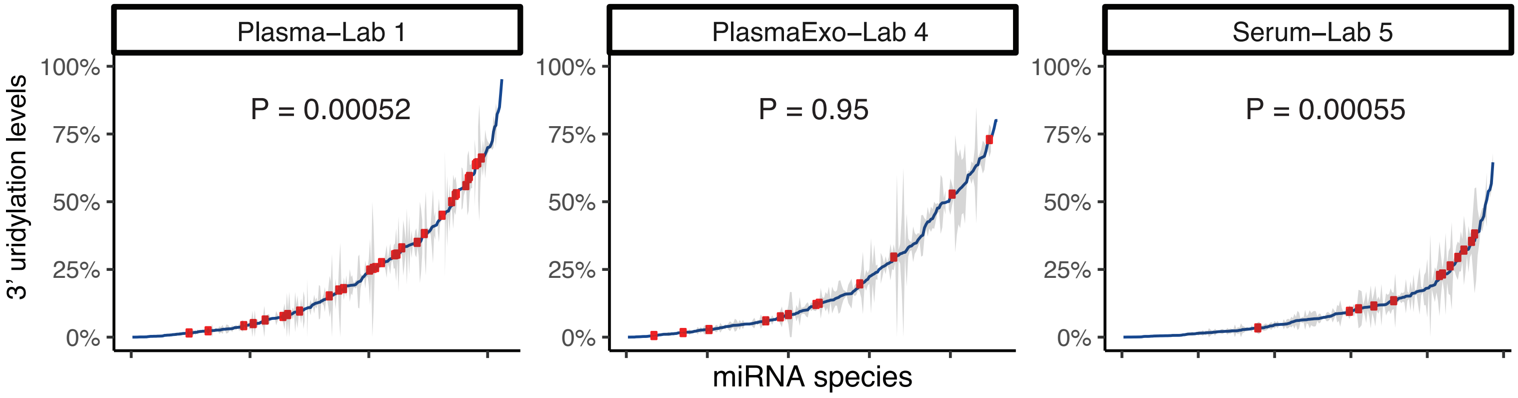


**Supplementary Fig. S5. Comparison of 3' uridylation levels in novel and known miRNAs.** 3' uridylation levels of miRNAs in each data set with at least 10 novel miRNAs expressed (read count ≥ 10). Blue curves represent the average 3' NTA levels and grey shades represent standard errors. Red dots represent novel miRNAs. P values were calculated via a two-tailed Wilcoxon ranked sum test.

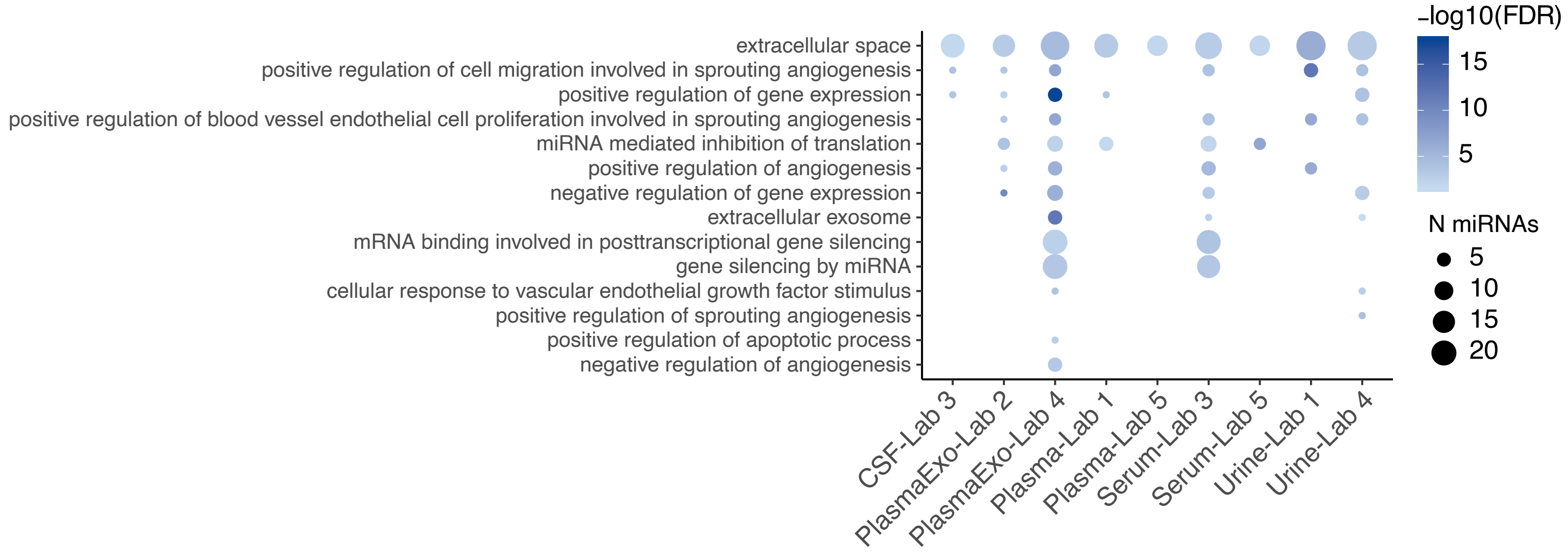


**Supplementary Fig. S6. Gene ontology enrichment of 3' uridylated miRNAs excluding let-7 family.** Gene ontology terms enriched among miRNAs, excluding the let-7 family, with an average 3' uridylation level ≥ 5% (FDR < 0.05, Methods). The size of the dots reflects the number of miRNAs in each GO term.

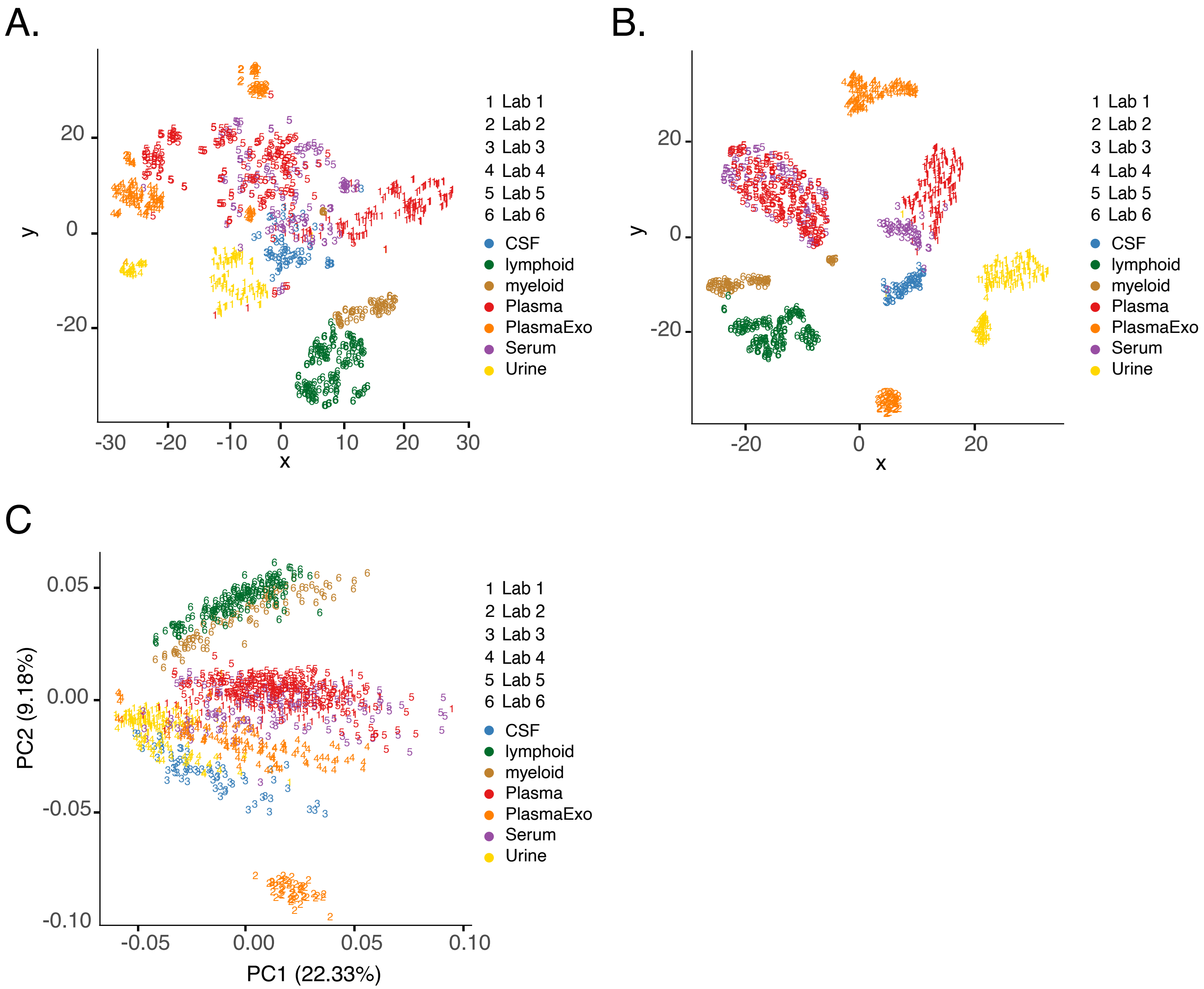


**Supplementary Fig. S7. Clustering samples from biofluids.**
(A) tSNE clustering of samples using miRNA 3' adenylation levels. miRNAs expressed with a minimum read count of 10 were included for this analysis (B) Similar to (A), but using normalized miRNA expression values (Methods). (C) Similar to (A), but PCA clustering of samples.
